# Microbial community dynamics in a traditional Swiss mountain cheese over 142 years of cheesemaking

**DOI:** 10.64898/2026.02.26.708305

**Authors:** Vincent Somerville, Marco Meola, Abigail Nunes-Richards, Johan Bengtsson-Palme, Judith Neukamm, Kerttu Majander, Marta Pla-Díaz, Meral Turgay, Sylvain Moineau, Monika Haueter, Hélène Berthoud, Ueli von Ah, Petra Lüdin, Verena J. Schuenemann, Noam Shani

## Abstract

The history of cheesemaking is deeply intertwined with the evolution of microbial communities, from spontaneous fermentation to modern, standardized practices. Despite centuries of refinement, the most profound shifts in cheese production occurred in the last century, driven by advances in microbiology and production technologies. These changes have shaped the bacterial and viral communities within cheese, yet the specific impacts remain underexplored.

Using shotgun metagenomics and 16S rRNA gene amplicon sequencing approaches, we examined microbial community changes in Raclette du Valais, a traditional Swiss cheese, using preserved cheese wheels from 1875 to 2017 from the same alpine dairy in Switzerland.

Our results reveal that significant shifts in microbial community composition coincide with changes in production practices. Notably, the oldest cheese harbored a distinct bacterial community, dominated by *Lactiplantibacillus paraplantarum*, *Streptococcus thermophilus*, *Pseudolactococcus laudensis*, and taxa commonly associated with the gut environment, indicative of spontaneous fermentation and the use of calf stomach for milk coagulation. Functionally, we can also track the rise and fall of antibiotic resistance genes mirroring their use. Furthermore, we found that domestication of lactic acid bacteria predates the studied period, and that bacteriophage genera detected in 1875 are representatives of those commonly found in modern cheesemaking. These findings highlight how microbial communities have adapted to changing production methods and how human intervention, through practices like antibiotic use in animal husbandry, has influenced these ecosystems in remote alpine cheesemaking.

**Significance Statement:** Cheesemaking relies on complex microbial ecosystems shaped by long-standing human practices, yet how these communities responded to the modernization of food production has remained largely unknown. By analyzing DNA preserved in historical cheese wheels from a single alpine dairy, we examine microbial community changes across a key technological transition. We show that modernization impacted bacterial composition and functional potential, that cheese microbiomes record the rise of agricultural antibiotic use, and that major cheese-associated phages and domesticated starter bacteria were already established over a century ago. These findings demonstrate that historical cheeses preserve long-term microbial records and offer a glimpse how changes in food production practices shape fermented-food microbiomes.

## Introduction

Human practices have played a major role in shaping both the use and the evolutionary trajectories of microbial communities involved in food fermentation. Initially, the surplus of food from the shift to a herding lifestyle was serendipitously preserved by spontaneous fermentation (1). Shortly after, the deliberate reuse of fermented products to initiate new fermentations emerged (2). In recent years, focus screening, selection and optimization of microbial cultures for food applications have regained momentum as fermented foods have experienced renewed popularity and interest (3).

Hardly any other fermented food product has undergone such diversification as cheese. Since the earliest evidence of cheesemaking around 5200 BCE (4), the 400 to over 1000 cheese varieties enjoyed today have evolved through millennia of adaptation to climate, preservation needs, milk sources, cultural traditions, and trade, yielding distinct yet reproducible tastes and textures (5, 6). Despite these ancient origins, cheesemaking has likely undergone its most profound transformations in the past century, driven by advances in microbiology and technology that enabled major increases in production, standardization, and food safety (Fig. 1) (7).

**Figure 1.**
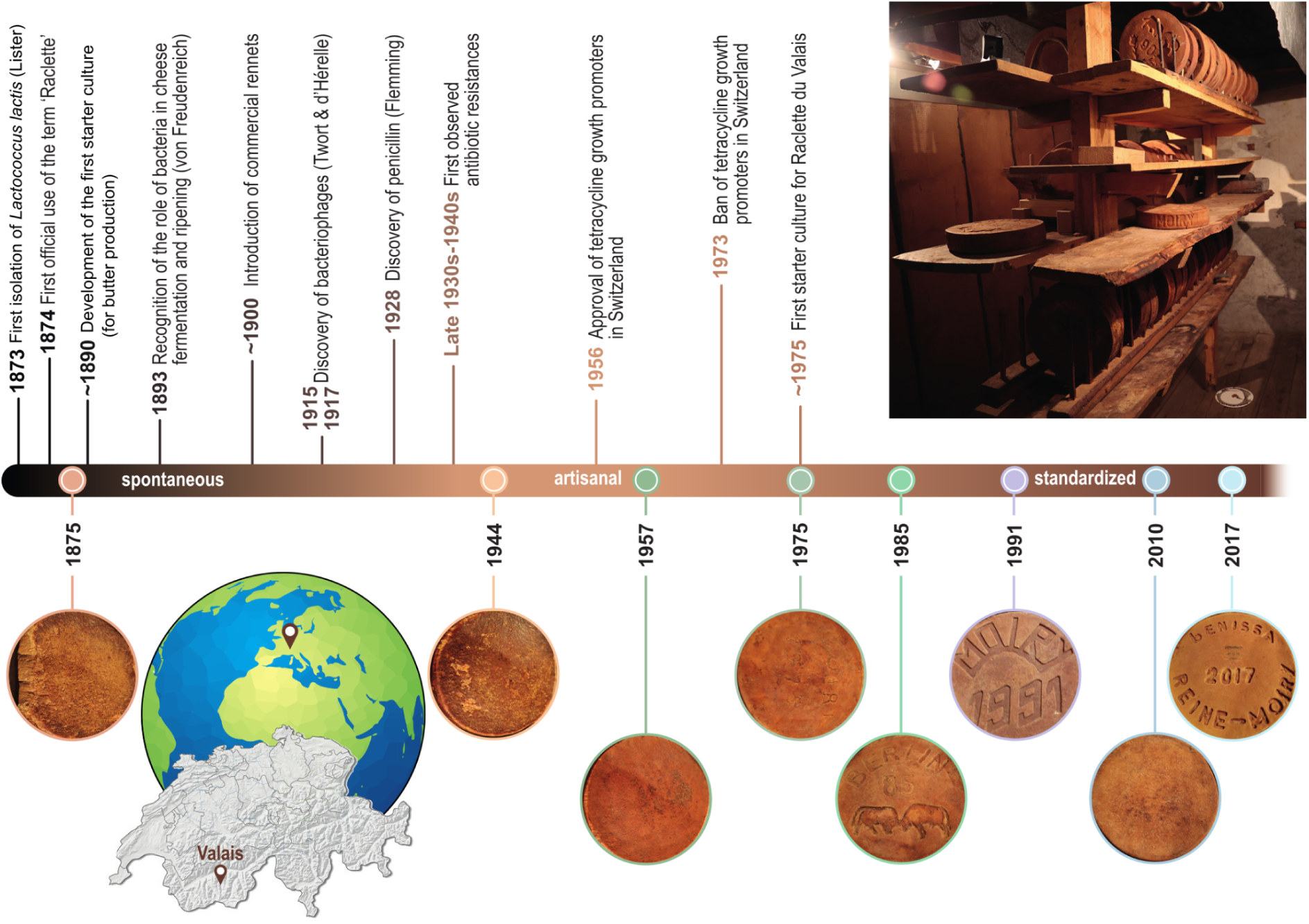
Timeline of the cheese samples investigated in this study and key technical and scientific events likely to have shaped cheese microbial communities. The globe on the left indicates the location of Switzerland and Grimentz, in the canton Valais, where the cheese wheels were sampled.

From a microbiological perspective, the last two centuries of cheesemaking can be divided into three partly overlapping periods, which emerged at different paces across regions. Until the turn of the 20th century (the ‘spontaneous period’), microbiological knowledge in cheesemaking was minimal and milk coagulation relied on spontaneous fermentation and/or on forms of back-slopping, the practice of reusing microbial communities from previous batches (8, 9). Despite this empirical approach, cheese was already recognized as a relatively standardized product, with cheese types displaying their specific characteristics (8). Reflecting this emerging standardization, *Lactococcus lactis,* the most common cheese starter bacterium used today, was first isolated in pure culture by Lister in 1873 (10). The following ‘artisanal’ period saw growing recognition of the microbial role and refinement of cheesemaking techniques (1), most notably through the utilization of natural starter cultures obtained through back-slopping or simply by incubating raw milk at room temperature, and the use of commercial rennet (11). This historical period extended until the adoption of commercial starter cultures, which occurred globally around the early-mid 20th century, though this transition occurred later locally (7). It also coincided with the discovery of antibiotics, which were widely and often excessively used or misused for human healthcare and animal husbandry in the second half of the 20th century. Beyond therapeutic applications, antibiotics were also used as growth promoters and for prophylactic interventions in livestock production, representing a major driver of antibiotic resistance development (12). Finally, the use of commercial starter cultures marked the beginning of the ‘standardized’ period. This innovation, together with increased mechanization and enhanced hygiene standards, enabled industrial-scale production. Technical improvements such as temperature control during cheesemaking further allowed for more standardized and reproducible processes. Hence, this period was also marked by growing knowledge of cheese microbiology as well as cheese chemistry and biochemistry and the advent of commercial fermentation-produced chymosin, which replaced the traditional rennet extracted from calf stomachs and the microbial rennets used after the 1960s (7, 13).

While the exact timing of transitions between these periods varied by region, cheese type, economic significance of cheese, and agricultural policies, their microbiological impact was likely far-reaching. Improvements in hygiene were associated with declining bacterial diversity in cheese (14). In parallel, antibiotics altered milk quality and microbial composition, at times even causing cheesemaking failures (15–17), while also contributing to the emergence and spread of antimicrobial resistance (17, 18). Finally, profound shifts in cheese bacterial community composition have probably been driven by transitions in milk inoculation practices: from spontaneous fermentation involving microorganisms from milk, the dairy environment, and calf rennets, to back-slopping and artisanal starter cultures, and ultimately to commercial, standardized starter cultures. These microbial transitions likely had cascading effects throughout the cheese microbiome, influencing not only bacterial diversity but also the composition and evolution of associated bacteriophage populations.

Despite major progress in understanding and controlling cheesemaking processes during the last century, lytic bacteriophages continue to pose a significant risk to consistent cheese production (19). Phages were co-discovered in 1915 and 1917, respectively by Frederick Twort and Félix d’Hérelle, with the first report of lactic acid bacteria (LAB) phages appearing in 1935, though these viruses were likely present since the earliest days of cheesemaking (20, 21). The phages found in modern cheese dairies, adapted to the narrow range of LAB used in starter cultures, likely differ substantially from those circulating during the spontaneous fermentation era. For example, the recently described *Piorkowskivirus* phage genus infecting *Streptococcus thermophilus* strains, another LAB frequently used in cheese production, appears to have emerged through recombination between a P335-type *L. lactis* phage and an unidentified *S. thermophilus* phage (22). This adaptive mosaicism event was likely favoured by the close ecological proximity of these bacterial species in milk and dairy environments, further intensified by the widespread use of starter cultures.

Although the three cheesemaking periods are relatively well defined technologically, assessing their impact on cheese bacterial and viral communities has long been hampered by the lack of historical cheeses spanning all production eras. Two approaches have therefore been used to study microbial change over time: (i) comparing genomes of contemporary microbial communities from different environments, and (ii) analyzing ancient samples. Modern genomes reveal population structure, but only ancient samples can show how microbial communities and genomes evolved. Several ancient cheese remnants have been discovered (23, 24), but they are isolated snapshots of early dairy practices and cannot capture the key microbial shifts of the last century.

Our study was based on a rare historical series of samples rooted in small alpine valleys of the canton Valais, Switzerland, where a transhumance-related funerary tradition, the ‘meal of the dead’, persisted into the early 20th century. In this custom, a cheese wheel acquired on a special occasion was kept until the owner’s death and served at his funeral (25, 26). As a remnant of this tradition, a few cheese wheels were still produced and kept throughout the late 19th and 20th centuries for commemorative purposes (27). Here, we obtained eight well-preserved cheese wheels, all produced at the same alpine pasture over 142 years (1875–2017). This unique chronological archive allowed us to trace microbial community dynamics and evolution across this critical phase of artisanal cheesemaking. The ‘cheese of the dead’ was a Raclette du Valais, a traditional semi-hard, smear-ripened cheese from the Swiss canton Valais. Made from raw cow’s milk, Raclette du Valais has been produced for centuries and remains in production today (28).

Here, we show that (i) ancient microbial DNA can be successfully recovered from historical cheese wheels, (ii) bacterial community compositions were shaped by cheese technological evolution, (iii) the impact of antibiotic use in veterinary medicine and animal husbandry reached even remote alpine production sites, (iv) domestication of LAB predates the studied period and (v) the most common bacteriophages in modern cheese dairies were already present 150 years ago.

## Results & Discussion

### DNA within the cheese matrix has remained well preserved for 142 years

To explore microbiological changes over more than a century, we extracted and sequenced DNA from eight cheese wheels produced between 1875 and 2017 (Fig. 1). Shotgun metagenomic sequencing yielded between 4.7 and 122 gigabases (Gb) per sample. The majority of reads were of prokaryotic origin (45–85%, Fig. 2A), with the remainder largely attributed to eukaryotes (mean = 33%) and phages (mean = 5%). Most eukaryotic reads originated from cows, with traces of sheep and goat DNA that were absent in the 2010 and 2017 samples (Fig. 2B), indicating that sheep or goat milk was still used at this particular Alpine dairy until three decades ago.

**Figure 2.**
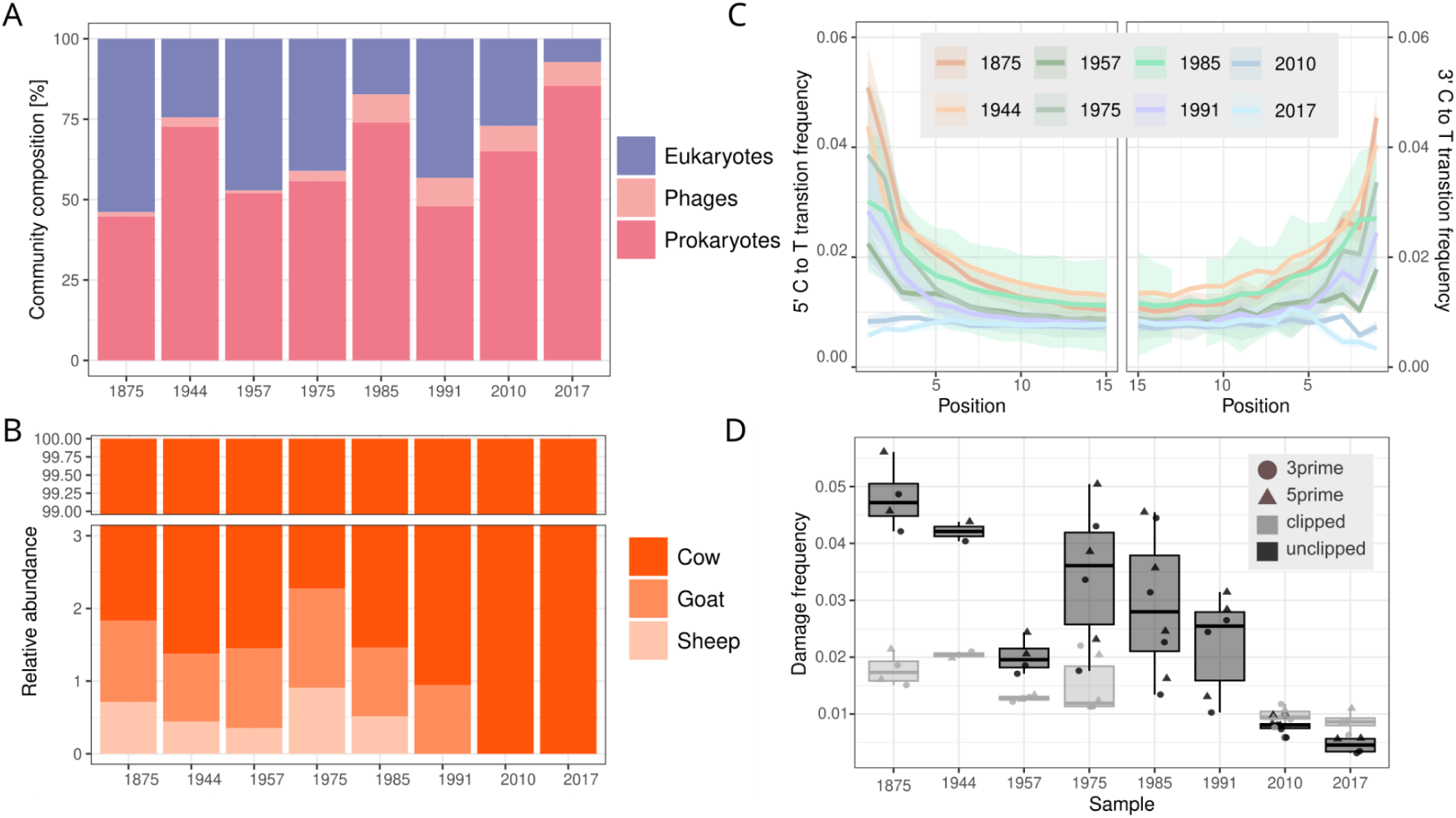
DNA sequencing and degradation over time. Shown are (A) the proportion of prokaryotic, eukaryotic, and phage reads, (B) the origin of eukaryotic reads, categorized as cow, sheep, or goat, (C) the damage frequency at the 5′ and 3′ ends of the reads, and (D) the changes in end damage frequencies over time.

Although milk is a known cryoprotectant (29), little is known about the long-term stability of DNA in processed milk stored at cellar temperature. To assess DNA quality and minimize artefacts, we examined nucleotide misincorporation patterns at the 3’ and 5’ ends of the DNA fragments (30). We focused on four key bacterial species typically present in Raclette du Valais cheese, *Lactococcus lactis*, *Lactococcus cremoris*, *S. thermophilus*, and *Lacticaseibacillus paracasei* (31) and considered the damage only if mapping coverage was higher than 10. Damage frequencies at both the 3’ and 5’ ends ranged from 0.003% to 0.05% and declined sharply within the first 6 bp (Fig. 2C). This degradation signal decreased significantly over time (0.028% per year; R^2^ = 0.51, p < 0.001, Fig. 2D). After clipping the first and last 6 bp, artefacts related to DNA diagenetic degradation were effectively removed (Fig. 2D).

Overall, the degradation gradient pattern expected in ancient DNA (aDNA) supports the hypothesis that the cheese samples analyzed contained aDNA with no modern DNA contamination. Our 1875 cheese samples exhibited DNA nucleotide misincorporation frequencies of 4-5% at the 3’ and 5’ ends, indicating relatively modest degradation for 150-year-old material. These values are comparable to those observed in oral microbiomes from the 16th-17th centuries (4-8%) and substantially lower than those from medieval samples (1000-2000 year-old oral microbiome samples: 20-30%) and a 5700-year-old oral microbiome (12.7-17.2%), suggesting that cheese represents a relatively stable matrix for long-term microbial DNA preservation (32–34). This opens new opportunities for high-resolution analyses of ancient food microbiomes, including functional, species-, and strain-level profiling. Recently, Liu et al. (2024) confirmed this potential by characterizing the microbiome of a 3500-year-old kefir cheese using shotgun sequencing and reconstructing the genome of *Lactobacillus kefiranofaciens* through targeted DNA capture followed by sequencing (24).

### Bacterial community shifts reflect 20th-century cheesemaking modernization

Cheesemaking typically relies on a specialized community of bacteria responsible for milk fermentation. To observe changes in the bacterial community composition, the 16S rRNA gene target sequencing was performed. Between 105 k and 1.4 M reads were obtained, resulting in 1326 different amplicon sequence variants (ASVs) from 305 different bacterial species. The species were categorized into four groups: ‘starter lactic acid bacteria’ (SLAB), ‘dairy bacteria’, ‘animal-associated bacteria’, and ‘other bacteria’ (Fig. 3A). Thirty-two species exceeded 0.1% median relative abundance (Supplementary Fig. 1). The cumulative relative abundance of subdominant species (< 0.1%) ranged from 0.6 to 7.5% of the community and declined over time in all four defined groups of species (Supplementary Fig. 2; linear model: *R^2^* = 0.49, *p* = 0.014). However, across all species, neither species richness (Supplementary Fig. 3; linear regression, *R^2^* = 0.0679, *p* = 0.2652) nor Shannon diversity (Supplementary Fig. 3; linear regression, *R^2^* = 0.1777, *p* = 0.164) declined significantly over time. This likely reflects functional redundancy among closely related species and the naturally high variability in dairy species diversity (31). In contrast, richness at the family level, which is likely a better proxy for broader functional diversity in dairy products, declined over time (Fig. 3B; linear regression, *R^2^* = 0.3179, *p* = 0.0663).

**Figure 3.**
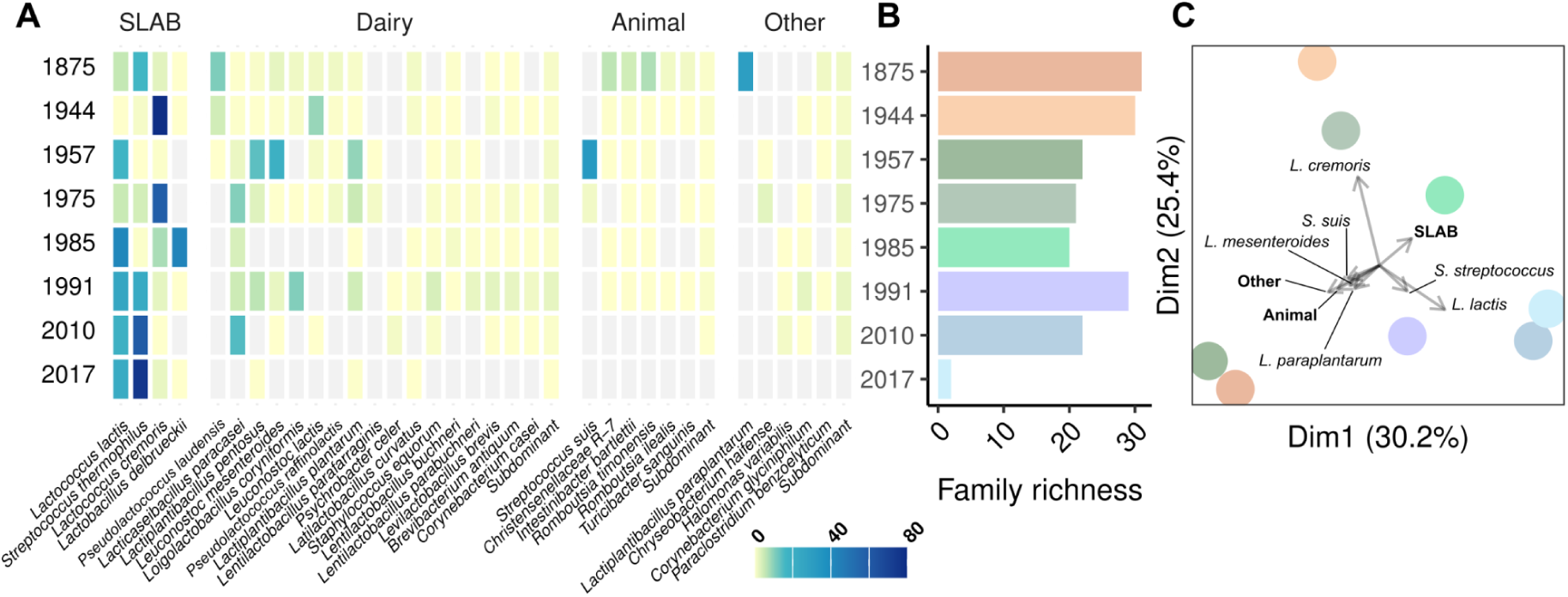
Temporal variations in the bacterial communities. A) relative abundance of the bacterial species over all samples grouped by starter lactic acid bacteria (SLAB), dairy bacteria (Dairy), animal-associated bacteria (Animal), or other bacteria (Other); B) Family-level richness; C) Principal Component Analysis (PCA) ordination biplot of the different samples based on relative abundance at species-level and categories. The colors of the objects refer to the year of cheese production and correspond to the colors of B).

Further, we assessed the temporal changes of the microbiota composition at the species level (Fig. 3A & C). The 1875 cheese sample was notably divergent, despite the presence of typical starter species. This was primarily due to (i) a comparatively higher relative abundance (21.1%) of strict animal-associated bacteria and (ii) high relative abundances of one dairy species, *Pseudolactococcus laudensis* (11.1%), as well as of the plant-associated *Lactiplantibacillus paraplantarum* (28.9%), all of which declined markedly in subsequent samples. As the animal-associated species did not belong to the typical indicators of fecal contamination (i.e., *Enterobacteriaceae* and *Enterococcaceae*) but were rather strict anaerobes typical of calves’ stomachs (35, 36), their presence likely resulted from the rennet extraction from a calf’s stomach using fresh whey, a common practice at the time (37) that disappeared with the advent of commercial rennets toward the end of the 19th century (38). *L. paraplantarum* could originate from several sources: traditional plant-based treatments for mastitis (39), the dairy environment and utensils, or a combination thereof. Notably, its high relative abundance suggests active proliferation during cheese maturation rather than mere persistence as a contaminant. This substantial growth indicates that *L. paraplantarum* likely played a significant functional role in the fermentation and ripening processes of this traditional cheese.

Three SLABs, *L. lactis*, *L. cremoris*, and *S. thermophilus,* were the most prevalent species, with their relative abundance significantly increasing over time, particularly from 1944 onward (Fig. 3A). The large dissimilarity between 1875 and 1944 reflects a profound shift in cheesemaking technology, transitioning from the ‘spontaneous’ to the ‘artisanal’ period. During the ‘artisanal’ phase, the mesophilic SLAB population, primarily composed of *L. lactis* and *L. cremoris*, clearly dominated over the thermophilic population, with *S. thermophilus* being nearly absent. This aligns with historical practices, where cheesemaking was initiated either with milk cultures obtained through natural fermentation at room temperature, akin to butter production (40), or with natural whey starter cultures (11). The latter method involved back-slopping, where the bacterial community recovered from a previous cheese production was used to inoculate the milk for subsequent production batches. Despite the close proximity of the 1957 and 1875 samples on the PCA plot (Fig. 3C), their bacterial communities differed markedly. The 1957 sample was dominated by *Streptococcus suis*, an animal pathogenic species absent in 1875 and detected only at very low relative abundances in all other samples. Conversely, *L. paraplantarum* was highly dominant in 1875 but absent in 1957. The high relative abundance of the mesophilic *L. lactis* in 1957 is consistent with the classification of this sample within the ‘artisanal’ period.

From 1985 onward, the thermophilic SLAB population significantly increased, signaling the onset of the “standardized” period, characterized by the use of commercial starter cultures, which resulted in reduced numbers of dominating bacterial species. Commercial cultures in use for Raclette du Valais production at that time combined mesophilic (*Lactococcus* spp.) and thermophilic (*Lactobacillus delbrueckii* subsp. *lactis* and *S. thermophilus*) strains. The detection of both lactococci and *L. delbrueckii* confirms commercial culture use in this alpine dairy. The 2017 sample contained the dominant species (*S. thermophilus*, *L. lactis*, *L. cremoris*) typically found in current cheeses of this type. However, non-starter LAB characteristic of Raclette du Valais, such as *L. paracasei*, were present only in low relative abundances, making this sample not fully representative of a typical modern Raclette du Valais (31).

These findings, together with historical records, illustrate the temporal evolution of cheesemaking practices from the late 19th century to the present day. These changes were driven by successive technological developments in cheesemaking: (i) the ‘spontaneous’ period, characterized by spontaneous fermentation and milk coagulation using calf stomachs, and reflected in the high relative abundances of species typically associated with the gut and plants; (ii) the ‘artisanal’ period, marked by the introduction of artisanal starter cultures produced through milk or whey fermentation at room temperature, combined with the use of commercial rennets, and associated with increased relative abundances of mesophilic LAB species; and (iii) the ‘standardized’ period, defined by the use of commercial starter cultures and rennet, and characterized by high relative abundances of a limited set of mesophilic and thermophilic LAB species.

### Conserved cheese-associated traits amid temporal fluctuations of ARGs

To explore functional changes in the cheese microbiome across time, we individually assembled and collectively binned the metagenomes of all samples. The total assembly length (range from 7.7 Mb – 31.3 Mb) decreased in more recent samples (Fig. 4A), likely reflecting the overall lower species diversity (Fig. 3). Most contigs were successfully binned into metagenome-assembled genomes (MAGs) across all time points (range from 48-88%, Fig. 4A). These bins represent a broad range of taxa and include 37 high-quality MAGs (completeness > 90%, contamination < 5%, Fig. 4B). Gene prediction within these MAGs revealed 96,070 different gene families. Consistent with the reduction in assembly size, gene richness also declined over time (Fig. 4C). This loss suggests a progressive narrowing of functional potential, likely related to the increasingly controlled conditions in the cheese dairy favoring mesophilic LAB and *S. thermophilus*. The 1875 sample was distinct regarding gene content (Supplementary Fig. 4), reflecting its unique bacterial community.

**Figure 4.**
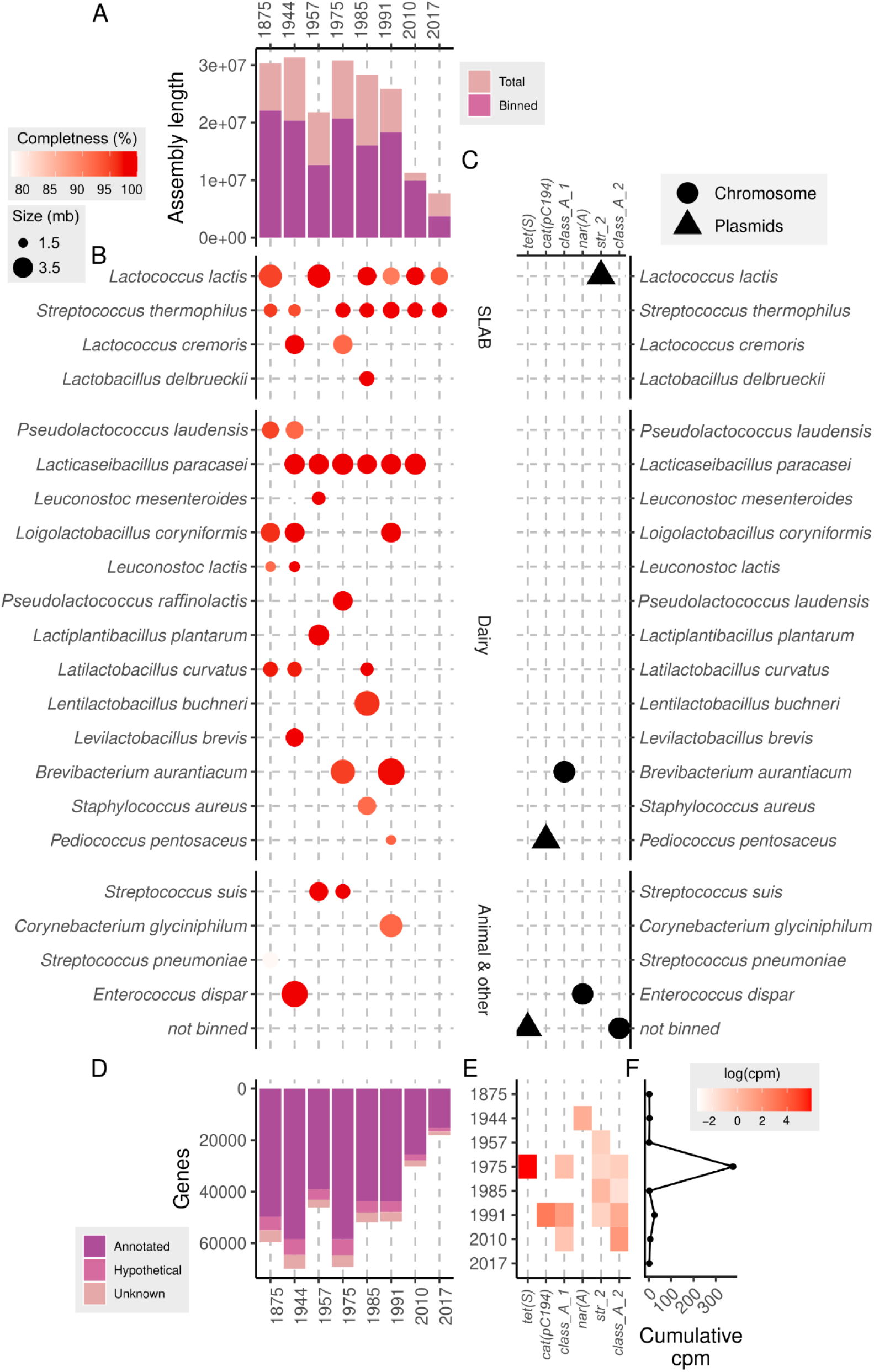
Functional diversity across samples. (A) Total assembly length for each sample, colored by binning status of contigs; (B) Metagenome-assembled genomes (MAGs), with color and size indicating respectively completeness and contig length; (C) Number of annotated genes per sample; (D) Genomic location of ResFinder-identified antibiotic-resistance genes (ARGs) within MAGs; (E) Normalized abundance (log(cpm)) of ARGs across samples; (F) Cumulative normalized ARGs counts per sample. cpm: copies per MAG.

The microbial functional gene content across all samples covered all core cheese-associated traits such as acidification (e.g., lactate dehydrogenase genes) and casein degradation (e.g., extracellular proteinases), which are primarily associated with SLAB (Supplementary Fig. 5) (41). The composition of cheese samples made it possible to track acquired antibiotic resistance genes (ARGs) across centuries. That allowed us to determine whether the spread of ARGs during the 20th century as a result of the widespread use of antibiotics in human, veterinary medicine, and animal husbandry was reflected in the cheese samples of this study. We thus hypothesized that ARGs detected using ResFinder (42) and fARGene (43) would appear after the discovery of antibiotics, that their number and diversity would increase as the 20th century progressed, and that they would persist over decades. While most samples contained few or no detectable ARGs, certain time points, particularly in the second half of the 20th century, exhibited sporadic surges in specific resistance determinants (Fig. 4D & E). As expected, MAGs from 1875 were devoid of acquired clinical ARGs. Interestingly, MAGs from 2017 were also free of such genes, possibly reflecting the effects of stricter regulatory measures implemented after 2012, when the European Food Safety Authority (EFSA) introduced microbiological cutoffs for clinically relevant antibiotics and issued recommendations to reduce ARG transmission through the food chain (44). In contrast, MAGs from 1944 to 2010 contained six clinical ARGs targeting different antibiotic classes (Fig. 4D & E): *tet(S)* (tetracycline resistance, tetracycline class), *cat* (chloramphenicol resistance, amphenicol class), a class *A1* β-lactamase (β-lactams class), *nar(A)* (narasin resistance, polyether ionophores class), *str_2* (streptomycin resistance, aminoglycosides class), and a class *A2* β-lactamase (β-lactams class). In the sample from 1944, only *nar(A)*, which confers resistance to narasin, was detected in the chromosome of *Enterococcus dispar*. Notably, polyether ionophores, the antibiotic class which narasin belongs to, were first isolated from *Streptomyces* sp. cultures in 1951 (45). Polyether ionophores were not introduced into agriculture to control coccidiosis in poultry until the 1970s and narasin was only characterized in 1978 (46, 47). The most abundant ARG in the MAGs, *tet(S)*, conferring resistance to tetracycline, was found exclusively in the sample from 1975. *tet(S)* genes have been previously identified in both Gram-positive and -negative bacteria, including *L. lactis* (48, 49). Our analysis also linked *tet(S)* in the 1975 sample to *L. lactis* based on contig assembly. Tetracyclines, first commercialized in 1948, gained rapid adoption in veterinary medicine due to their broad-spectrum activity and utility as growth promoters and prophylactics (17, 50). In Switzerland, they were used in veterinary medicine from the 1950s and authorized for growth promotion in livestock between 1956 and 1973 (51). The high abundance of *tet(S)* suggests that the strain harboring it was dominant, although its origin, milk or starter culture, remains unclear.

A gene conferring resistance to aminoglycosides, *str_2*, appeared in four consecutive samples, from 1957 to 1991. Aminoglycosides, introduced in 1944, were widely used in agriculture, often alongside β-lactams (52). Two genes predicted to confer resistance to β-lactams were detected in samples from 1975 to 2010. One (class_A1) shared 97% with a gene from *Brevibacterium aurantiacum* strain SMQ-1421, and the other (class_A2) showed 100% identity with a gene in *Staphylococcus* sp. OJ82. Finally, an ARG (*cat*) conferring resistance to chloramphenicol was identified in the sample from 1991. Although chloramphenicol was widely used to treat bovine infections since the 1940s, it was banned in the EU for food-producing animals in 1994 (53).

The temporal pattern described here underscores the impact of antibiotic use in agriculture on the development and dissemination of antibiotic resistance genes (ARGs) in cheese production. Before the 1950s, antibiotics were not used in animal husbandry in Switzerland; accordingly, the samples from 1875 and 1944 contained either no ARGs or only one at low abundance. A marked increase in ARG prevalence and abundance was observed after the introduction of antibiotics as growth promoters for livestock. The most abundant ARG, *tet(S)*, detected in the 1975 sample, likely reflects the use of tetracyclines as growth promoters from 1955 until their ban in 1973. The widespread use of antibiotics during this period likely exerted substantial selective pressure that drove the dissemination of ARGs even to remote environments such as alpine pastures, where growth promoters were not used for dairy cattle, with ARGs persisting shortly after their official withdrawal. Finally, the introduction of recommendations and stricter measures in food and feed production coincides with the disappearance of ARGs in the most recent sample. The relatively rapid disappearance of *tet(S)* (within less than two decades after the ban on tetracycline growth promoters) contradicts our hypothesis of long-term ARG persistence and suggests that appropriate regulatory measures can effectively mitigate ARG dissemination and persistence.

### Major LAB domestication occurred prior to the period analyzed

Despite changes in the overall microbial community composition, the bacterial species responsible for milk acidification are consistently reused in cheesemaking, namely as SLAB. As a consequence, these starter bacteria bear genomic imprints of domestication (54). While this process has been going on for millennia, we wanted to assess domestication imprints accumulating in the timescale of these samples. We examined key features at three levels: community, genome, and gene.

To assess whether industrialisation has reduced functional gene diversity in the bacterial communities, particularly in cheese starter bacteria, analogous to the loss of diversity observed in crop monocultures (55), we quantified temporal changes in core-genome nucleotide diversity (SNPs per 100 bp) in *S. thermophilus* and *L. lactis*. Overall, both species exhibited low nucleotide diversity, ranging between 0.005 and 0.1 SNPs per 100 bp. Low nucleotide diversity has previously been identified as a hallmark of traditional cheese starter cultures as well (41). However, no distinct decline in nucleotide diversity over the studied timeframe was observed (Fig. 5A). We found no strong evidence that these species occupy different niches in the cheesemaking process, which may explain the generally limited standing strain variation.

**Figure 5.**
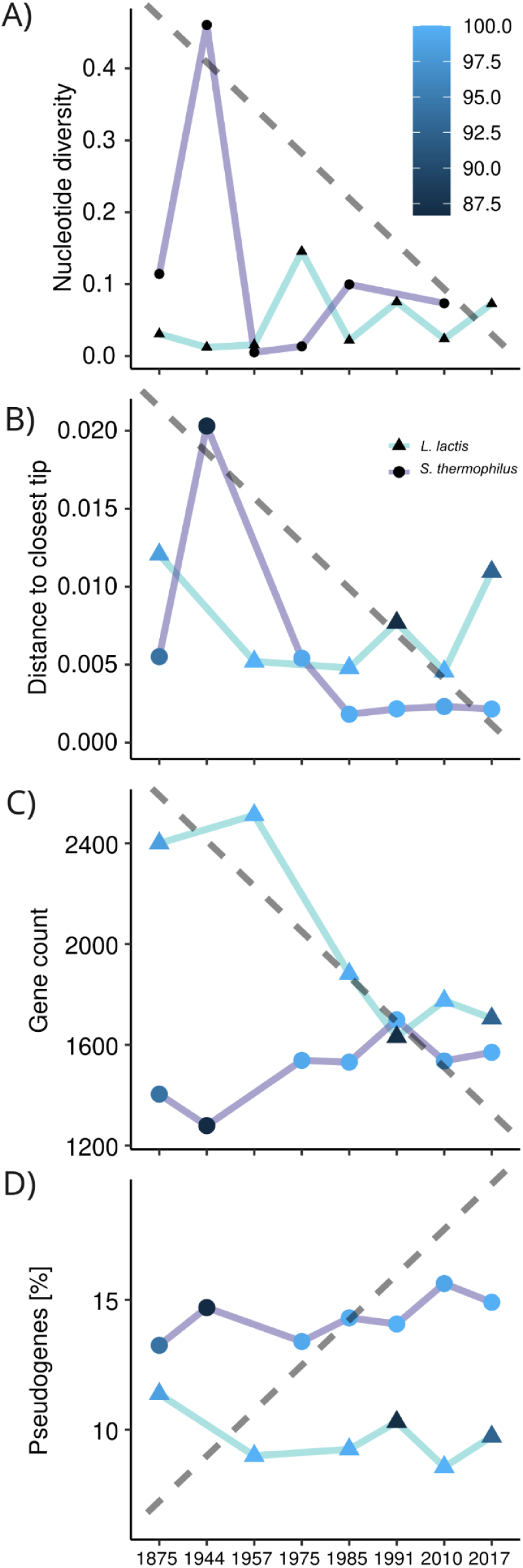
Four aspects of domestication (from top to bottom). Expectations under a domestication scenario are illustrated as dashed grey lines. A) Nucleotide diversity of the different samples for *L. lactis* and *S. thermophilus*.; B) Distance to the closest non-MAG tip for the different MAGs; C) Pseudogene and D) gene counts of the *L. lactis* and *S. thermophilus* MAGs.

Historically, dominant strains may have differed from those present today, analogous to the replacement of regionally adapted crop varieties by a few global breeds (56). To test whether contemporary strains are more closely related to recent communities, we constructed a phylogeny of MAGs and reference isolates and quantified each MAG’s distance to its nearest phylogenetic neighbour (Fig. 5B and Supplementary Fig. 6/7). In general, closely related reference strains were identified for all MAGs, with distances ranging from 0.002 to 0.01 SNPs per 100 bp. However, these phylogenetic distances remained stable over time, suggesting that historical strains have remained as similar to their closest relatives as more modern strains are.

Although we detected only single dominant strains in the historical samples, these species are known to have undergone gradual genetic erosion due to long-term, intensive use in cheese production (54). To test whether such erosion is also detectable within the timeframe covered by our samples, we compared gene and pseudogene counts across MAGs. Complete genomes of *S. thermophilus* and *L. lactis* typically contain relatively low gene counts (e.g., *L. lactis*: 2055 ± 374; *S. thermophilus*: 1546 ± 94) but high pseudogene counts (9.58 ± 1.09; 14.3 ± 0.905, respectively). Our MAGs showed slightly fewer genes and pseudogenes (Supplementary Fig. 8), likely reflecting incomplete assemblies (57). However, we found no evidence of temporal genome reduction (Fig. 5).

Overall, we detected genomic signatures consistent with domestication at the community, genome, and gene levels. However, the lack of temporal trends in our dataset suggests that most domestication likely predates the time window captured here. This aligns with genomic evidence indicating that the main dairy-associated lineages of LAB originated 2000-8000 years ago, with most diversification and adaptation occurring during that time (54). Given these long evolutionary timescales, the absence of measurable domestication signatures within the recent historical window examined here was not surprising; yet, to our knowledge, this lack of short-term evolutionary change has not yet been shown. Thus, the shift toward defined and repeatedly reused starter cultures likely imposed strong selection of specific strains rather than continued evolution within them, resulting in little measurable change over the historical timescale examined here.

### The worries are the same: overall stable phage communities

Phages remain a major risk in cheesemaking today, capable of causing production failures or quality issues by infecting starter bacteria such as *S. thermophilus*, *L. cremoris*, and *L. lactis* (58). To evaluate whether phage pressures have changed over the historical period covered by our samples, we first characterized the phage communities by identifying phage contigs in the assemblies with geNomad (59). In total, 68 distinct phages (≥ 99% identity and ≥ 85 % coverage) were identified. While 24 phages (41%) were unique to individual samples, 8 phages (14%) were shared between 5 to 8 samples (Supplementary Fig. 9). Notably, the damage profiles of these contigs exhibited the same progressive reduction with sample age (Supplementary Fig. 10) as observed for bacterial contigs (Fig. 2C), confirming they represent genuinely ancient phages rather than modern contaminants. Samples from consecutive timepoints were more similar in their phage community composition (Supplementary Fig. 11). Phage richness did not decline significantly over time (linear regression, p-value=0.159, r^2^=0.301, Fig. 6A), and viral load showed only a modest increase once adjusted for contig coverage (linear regression, p-value=0.08, r^2^=0.42, Fig. 6B). Together, these results indicate that phage communities have remained relatively stable in these cheeses across the examined historical period.

**Figure 6.**
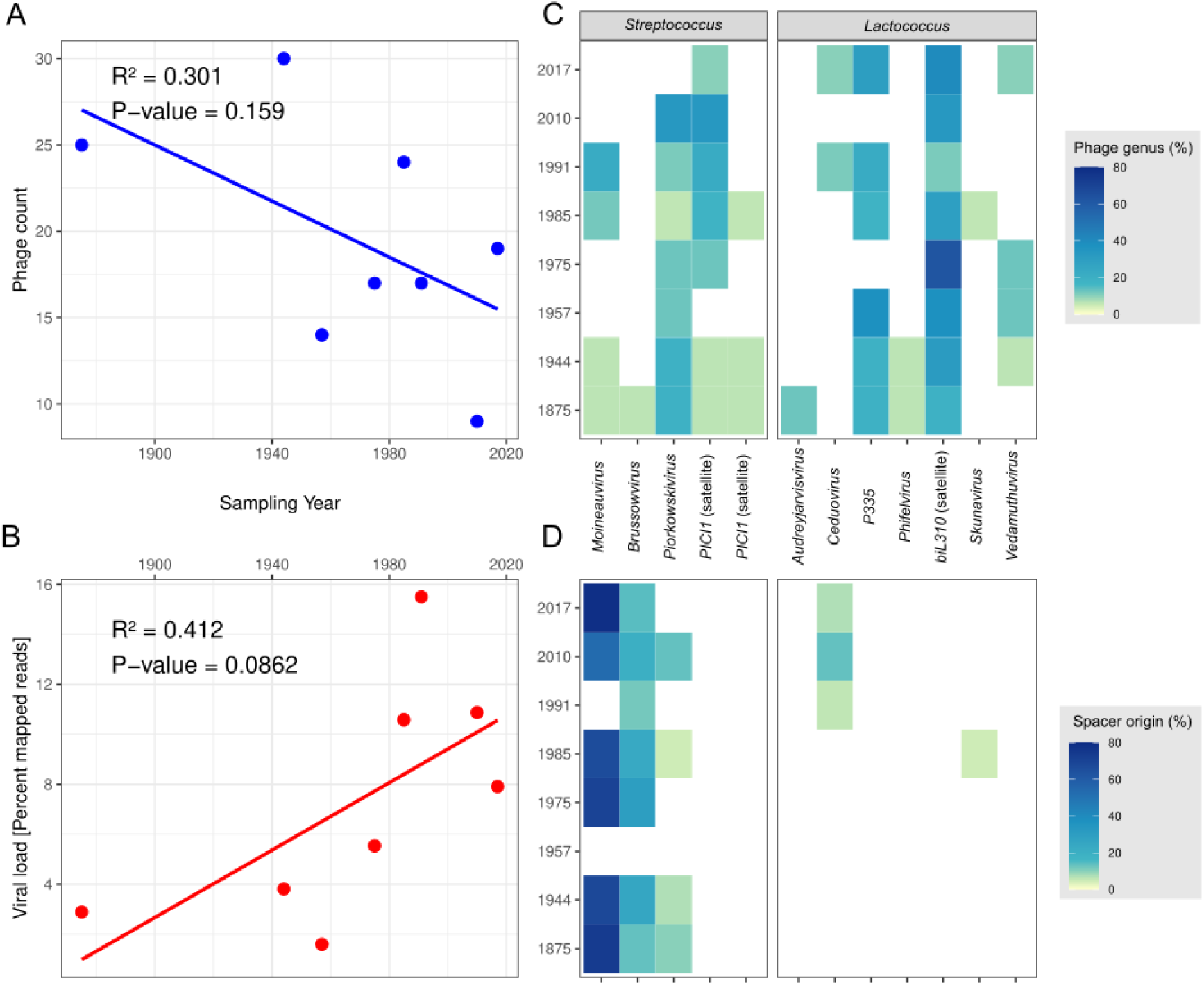
Phage diversity plot. Phage count (A) and viral load (B) over time with the linear regression illustrated as a line; C) Fraction of contigs coming from the different viral genera (and satellite phages) including only the clustered contigs; D) Fraction of mapped CRISPR spacers colored according to viral genera.

To determine whether this stability also extended to phage identity and host range, we then clustered and identified the phage contigs using vContact2 (60) and iPHoP (61). Approximately 21% of the phage contigs were associated with streptococcal, 38% with lactococcal and 2% with lactobacilli hosts (Fig. 6C). While the remaining 40% did not cluster with any known phages, they were also shorter than 5000 bp, indicating likely incomplete assemblies rather than novel phage groups. Consistent with the results above, we did not observe clear temporal shifts of specific phage genera (two-way ANOVA, p > 0.05), suggesting that the various genera overall coexisted consistently over time. This includes *Piorkowskivirus*, a hybrid lineage between an *S. thermophilus* and an *L. lactis* phage considered increasingly common in modern dairy environments (22), which was already consistently present in our dataset.

Given the importance of lactococcal phages in current cheese production, we examined how the three major contemporary taxonomic groups (*Skunavirus*, *Ceduovirus*, and P335) were represented over time. Members of the lactococcal P335 phage group were detected in nearly all cheese samples, in the form of prophages integrated into bacterial genomes.In contrast, the strictly lytic lactococcal phages belonging to the *Skunavirus* and *Ceduovirus* viral genera were detected only from 1985 onward, with *Skunavirus* present only in 1985 and *Ceduovirus* appearing in the 1991-2017 samples. Although modern cheesemaking environments are typically dominated by *Skunavirus* (62, 63), our results depict a slightly different situation in this specific cheese dairy, where *Ceduovirus* became more prominent in the later decades. Nevertheless, the fact that both these phage genera were not detected in our metagenomic data does not imply that they were absent. Rather, they may have been present at sufficiently low titers to escape detection by metagenomic approaches. Such low abundance would be consistent with the absence of production defects, as high titers of these lytic phages are typically associated with production failures.

To further investigate changes in *S. thermophilus* phages over time, we analyzed phage-targeting CRISPR spacers. All known strains of *S. thermophilus* carry CRISPR-Cas systems (64, 65). Notably, fragments of lactococcal phage DNA were detected in CRISPR-Cas systems, although *L. lactis* and *L. cremoris* rarely possess this anti-phage defense system (66). Across samples, we identified 1179 *S. thermophilus* CRISPR spacers distributed across three prevalent CRISPR arrays (2 type II-A and 1 type III-A), averaging 182 spacers per sample (SD = 113; Fig. 6D). Most spacers (85%) were unique to individual samples (Supplementary Fig. 12). When mapped to known phages from IMG/VR and our assembled phages, 14% of the spacers matched identifiable targets, predominantly streptococcal phages belonging to the *Moineauvirus* (69%) and *Brussowvirus* (20%) viral genera, which are also the two most predominantly found in the dairy industry today. Generally, spacer mapping is below 50% (65). With the exception of 1957, where a single *S. thermophilus* spacer was recovered due to *S. suis* dominance, no significant pattern emerged regarding targeted phage genera. Thus, CRISPR evidence supports the same long-term stability observed at the level of phage abundance and taxonomy.

Overall, these results indicate that phage communities and their interactions with host bacteria have remained relatively stable over the past 150 years. Both metagenomic evidence, including phage abundance and identity, and CRISPR analyses reveal no signs of major shift in phage prevalence or infectivity within this cheese production system.

## Conclusion

To understand the anthropogenic impact on the microbes in fermented foods, we studied the microbial dynamics in cheese wheels from the same type and geographic location over the past three centuries, a period marked by major scientific, technological, and industrial transformations in cheesemaking. Exploiting a unique regional tradition, where cheese wheels were kept as family heirlooms, we revealed major shifts and a general decline in taxonomic and functional diversity. Notably, both domestication and phage diversity predated the transitions in production practices and the adoption of defined industrial starter cultures.

Phylogenetic inference offers a powerful window into the historical gene pool, but it provides only a coarse view of past microbial diversity. Resolving how microbiomes truly varied through time requires direct analysis of historical and archaeological specimens, despite the inherent limitations of such archives, including restricted sample availability, lack of biological replicates, and the possibility of sample-specific anomalies. For example, the 1957 cheese was dominated by *Streptococcus suis*, whereas the 1985 cheese reflected the introduction of a thermophilic starter culture, evident from the unusually high abundance of *Lactobacillus delbrueckii*. Despite these constraints, museum collections, historical archives, and other preserved materials remain indispensable for reconstructing the evolutionary and ecological dynamics of fermented foods. Beyond changes in community composition, such archives uniquely capture shifts in the functional gene pool that cannot be inferred from phylogeny alone. In particular, they provide a rare opportunity to trace the emergence, spread, and decline of antimicrobial resistance genes in these microbiomes across the twentieth century, a period marked by the rapid expansion of antibiotic use in agriculture followed by increasing regulation (17). The temporal patterns observed in our dataset indicated that AMR gene presence closely tracked these broader regulatory and technological changes. While limited to a single cheese series, this pattern suggests fermented foods could serve as valuable recorders of anthropogenic selection pressures acting on microbial communities.

In contrast to the patterns observed for ARGs, bacteriophages were detected at all time points, with no strong evidence for major temporal shifts in overall phage genera or abundance across the dataset. Their consistent presence possibly indicates that phages were a persistent component of cheesemaking systems throughout the study period, despite substantial changes in bacterial community composition and production practices.

Across the full timeline, we observed a gradual reduction in overall functional diversity, likely driven by production standardization, improved hygiene, and widespread adoption of defined starter cultures. Core functions essential for cheesemaking, such as acidification and casein degradation, remained stable, yet the loss of broader metabolic capabilities may influence flavor complexity, nutritional composition (e.g., via shifts in microbial protein profiles), and ecosystem resilience (67, 68). Although sensory and biochemical effects were not measured here, recent advances in metagenomics and experimental reconstruction (69, 70) now enable the recreation of historical or synthetic cheese communities. Building on our metagenomic dataset, future work could attempt to reproduce cheeses resembling those produced 150 years ago, test whether similar trends occur in other cheese types, and experimentally re-establish historical production conditions to further clarify how human innovation has shaped, and continues to shape, the microbial ecosystems of fermented foods.

## Materials and Methods

### Cheese samples

The cheese wheels investigated in this study were produced at the same alpine dairy in the canton of Valais, Switzerland (alpine pasture of Moiry, former pasture of Torrent). This is the highest cheese dairy in Switzerland (2497 m ASL, GPS coordinates 46.131, 7.557). The only exception was the cheese wheel from 1944, manufactured on the alpine dairy of Châteaupré (J.-J. Zufferey, personal communication) at ca. 2250 m ASL, which was subsequently flooded as the Moiry dam was commissioned in 1958. This alp pasture was located in the same valley as the dairy of Moiry.The cheese wheels were preserved in the cellar of the Zufferey family, in Grimentz (Switzerland). Samples were selected based on having a known production date between 1875 and the present. The selection comprised 8 cheese wheels from the years 1875, 1944, 1957, 1975, 1985, 1991, 2010, and 2017.

### Sample collection and preparation

Samples of the 1875 cheese were taken using a sterilised scalpel in a dedicated ancient DNA clean laboratory at the Institute of Evolutionary Medicine, University of Zurich.

For the cheeses 1944, 1957, 1975, 1985, 1991 and 2017 two samples were collected. The first 1 cm (“plug”) was not included, the sample .1 was 1-4 cm and .2 was 4-8 cm into the cheese wheel. Sample 2010.1 included the plug and 2010.2 was 4-6 cm in depth. Samples from the 2017 cheese were taken using a sterile biopsy needle (BMN “J” Type 8Gx150mm, HS Hospital Services SpA, Italy) in a dedicated modern DNA extraction laboratory at the Institute of Evolutionary Medicine, University of Zurich. All other samples were taken using a sterile biopsy needle (ibid) *in situ* in the cheese cellar, Grimentz, Switzerland. All samples were stored at 4 °C after sampling.

### DNA extraction

For the pre-1970 samples, DNA extractions and sequencing library preparations were performed in the ancient DNA clean laboratory at the Institute of Evolutionary Medicine, University of Zurich (71) following standard anti-contamination protocols (72–74) with parallel non-template controls. For the post-1970 samples, the same anti-contamination procedures were enforced in a dedicated modern DNA extraction laboratory at the Institute of Evolutionary Medicine, University of Zurich. Approximately 100-150 mg of cheese were homogenized using a sterile 200 µl pipette tip and incubated at 55 °C and 600 rpm for 4 hours, then overnight at room temperature at 600 rpm in 1.25 L of extraction buffer (160 mM ethylenediaminetetraacetic acid, 2% sodium dodecyl sulfate, 4 µg/µl proteinase K), A total of 1 ml was transferred to 15 ml of binding buffer (5 M GuHCL, 40% Isopropanol, 400 µl sodium acetate 3 M; UV light treated before use). The mixture was then transferred to a high pure extender assembly (Roche Applied Science, UV treated) modified with a MinElute column (QIAGEN), centrifuged until all the sample solution was passed through, and washed twice with 700 μl PE buffer (QIAGEN) before elution in 100 μl of TET (10 mM Tris-HCl, 1 mM EDTA, 0.05% Tween-20, pH 8.0).

### Shotgun metagenomic sequencing

Metagenomic sequencing libraries were generated from 10 μl of extract or extraction blank following (75, 76) with modifications. The re-amplification step was performed with 1 unit Herculase II DNA polymerase (Agilent), 5× Herculase II reaction mix, 0.3 μM primers IS5 and IS6 (75), and 4-7 μl library template with the following thermal profile: initial denaturation at 95 °C for 2 min, 10 to 25 cycles of denaturation at 95 °C for 30 sec, annealing at 60°C for 30 sec, and elongation at 72 °C for 30 sec, followed by a final elongation at 72 °C for 5 min. Libraries were purified with MinElute spin columns following the manufacturer’s instructions. Quantitative PCR (qPCR) analysis on an Agilent 2200 TapeStation were used to assess the quality and concentration of the libraries. Libraries were pooled and sequenced on the illumina platforms NextSeq500 2×75 bp and NovaSeq 2x150 bp at the Functional Genomics Center Zurich (Switzerland). The raw reads were mapped, quantified, and removed against reference genomes of the dairy species *Bos taurus* mt genome NC_006853.1 and whole genome GCF_002263795.1_ARS-UCD1.2, *Capra hircus* mt genome NC_005044.2 and whole genome NC_030808.1, *Ovis aries* whole genome GCF_000298735.2, *Sus scrofa* mt genome KX094894.1 and whole genome GCF_000003025.6 and the human genome with kneaddata (v0.7.7-alpha) (77). Additionally all reads were clipped by 6bp to remove potential age damage (Fig. 2C).

### Metagenomics data processing

Shotgun metagenomic reads were assembled, binned, annotated, and quantified with ATLAS v.2.5.0 (78) using default parameters and kmers of 21, 33, and 55 for assembly with spades v.3.13.1, MaxBin v.2.2.7 (79), MetaBAT v.2.14 using minContig 1500, minCV 1.0, minCVSum 1.0, maxP 95%, minS 60, and maxEdges 500. Other tools used in ATLAS were: diamond v0.9.29.130, blastp 2.9.0+, Prodigal v.2.6.3, ruby v.2.6.5p114. Quality of the MAGs was assessed with checkM v.1.1.1 (80) and BUSCO v.5.8.0 (81).

### DNA damage analysis

The raw sequencing data were mapped against the complete reference genomes of *L. lactis* (RefSeq ID: NC_002662.1), *L. cremoris* (RefSeq NZ_BCVK01000000) *S. thermophilus* (RefSeq ID: NC_017581.1) and L. paracasesi (RefSeq ID: NZ_CP007122). The mapping and further processing was conducted as described above using the EAGER pipeline version 1.92.55 (82). In brief, the DNA fragments were mapped using BWA aln (83) with a mapping quality of 25 and a maximum edit distance of n = 0.1. All other parameters were used as default by EAGER (82). Duplicates were removed with MarkDuplicates (https://broadinstitute.github.io/picard) and Qualimap version 2.2.1 (84) was employed to evaluate the mapping. Finally, DamageProfiler version 1.0 (85) was used to investigate the damage patterns.

### 16S rRNA-based bacterial community analyses

#### 1 16S rRNA gene amplicon sequencing

Amplicon sequencing of the 16S rRNA gene was performed as described previously (31) with some modifications. Briefly, libraries were prepared by PCR amplification of the V1-V2 region of the 16S rRNA gene using primers 8F (5′-AGAGTTTGATCMTGGCTCAG-3′) and 355R (5′-GCWGCCTCCCGTAGGAGT-3′) with 35 to 40 cycles at 94°C for 30 s, 55°C for 30 s, and 68 °C for 30 s after an initial denaturation step at 94 °C for 2 min. The amplicons were sequenced on an Ion S5™ System with an Ion530 Chip (Thermo Fisher Scientific, Waltham, MA, USA).

#### 2. Determination of the bacterial composition

The single-end (SE) raw reads (mean length = 310 bp; sd = 32 bp) were primer trimmed and quality filtered (maxEE = 15, truncQ = 6, maxN = 0, n = 1e+06, minLen = 100, max-Len = 460) in DADA2 (86). Amplicon sequence variances (ASVs) were obtained in DADA2 with the parameter POOL = “pseudo”. Taxonomic annotation was performed using DAIRYdb v1.2.5 (87) with IDTAXA (88). ASVs assigned to *L. lactis* were further resolved to *L. lactis*, *cremoris* or *allomyrinae* by blastn v.2.13.0+ and the command: blastn -query [query_file.fasta] -db [16S_ribosomal_RNA.fasta] -out [output_file.txt] -outfmt “6 qseqid sseqid pident length mismatch gapopen qstart qend sstart send evalue bitscore salltitles” -num_alignments 5.

The bacterial communities were analyzed at the species level. Multivariate statistical analyses were performed using RStudio Pro v.1.4.1103-4 (RStudio Team (2020). RStudio: Integrated Development for R. RStudio, PBC, Boston, MA URL http://www.rstudio.com/) with the R software v.4.0.3 (R Core Team (2021). R: A language and environment for statistical computing. R Foundation for Statistical Computing, Vienna, Austria. https://www.R-project.org/) and the Phyloseq package v.1.42.0 (89).

#### 3. Analysis of changes in the bacterial community structures

The barchart was generated with normalized compositional samples. For clarity reasons, all species with abundances lower than 1% in all samples were removed. The richness and diversity plot was performed using the *plot_richness* function in Phyloseq with the not normalized counts at species level.

The PCA biplot was performed using the normalized species abundance table. The values were Hellinger-transformed using the decostand function in the vegan package v.2.6-4. The Principal Component Analysis (PCA) was calculated with the *prcomp* function in stats package with parameter scale=FALSE. The PCA biplot was visualized using the function *fviz_pca_biplot* from the factoextra package v.1.0.7.

### Analysis of the functional composition

The functional composition analysis was based on outputs generated by the ATLAS pipeline (v.2.5.0). We quantified the number of SPAdes-assembled contigs assigned to each DASTool bin to assess genome-resolved functional potential. We also enumerated gene families detected in each sample and recorded the species and normalised abundance (counts per million). Plasmid contigs were identified using geNomad (59). Antibiotic resistance genes were annotated using Usearch (90) against the ResFinder database (42) using a 70% identity and 80% target coverage cutoff. In addition, fARGene (43) was used to identify potentially novel ARGs.

### Genomic evolution analysis of key species

To quantify nucleotide diversity in *L. lactis* and *S. thermophilus*, quality-filtered reads were mapped to a reference collection of SLAB species genomes using BWA-MEM v.0.7.17 (91). Species-level mappings were included only when the relative abundance of the species exceeded 0.01%. Single-nucleotide polymorphisms (SNPs) were identified with FreeBayes (92), and we retained only positions with >10× coverage, located within the core genome, and exhibiting an allele frequency between 0.2 and 0.8. For phylogenetic analysis, we downloaded all available RefSeq proteomes for *L. lactis* and *S. thermophilus*. A core-proteome phylogeny was inferred using OrthoFinder v2.5.5 (93) and phyloplan (94) and visualised with iTOL (95). Distances between MAGs and their nearest phylogenetic neighbors were calculated in R using the find_nearest_tips function from the castor package v.1.7.4. To further characterize genome integrity, gene sequences were extracted with Bakta v1.9.2 (96), and pseudogene counts were identified using PseudoFinder v.1.1.0 (97).

### Analysis of the bacteriophage communities

Bacteriophage contigs were identified from the ATLAS assemblies using geNomad v.1.7.4 (59). Only contigs longer than 1 kb were retained. Contigs were de-replicated at 99% nucleotide identity and 85% alignment coverage using a combination of BLASTn v.2.14.1 (98), aniclust v.1.0.2, and seqkit v.2.6.1 (99). Following de-replication, only phage clusters with a total length of at least 5 kb were retained for downstream analyses. Genome completeness was assessed using CheckV v.1.0.1 (100), bacterial host prediction was performed with iPHoP v.1.3.3 (61), and phage lifestyle was inferred using BACPHLIP v.0.9.6 (101). Phage prevalence and abundance were estimated by mapping metagenomic reads to the phage reference collection using BWA-MEM v.0.7.17 (91), and coverage was calculated with CoverM v.0.6.1 (102). Phage genomes were annotated with Pharokka v.1.7.2 (103). Viral clustering and taxonomic assignment were performed using vContact2 v.0.11.3 (60) with the RefSeq viral reference database (downloaded 20241201). CRISPR spacers from *S. thermophilus* were extracted directly from metagenomic raw reads as previously described (65). Briefly, conserved CRISPR repeat sequences were used to identify and excise CRISPR arrays from raw reads using Cutadapt v.4.6 (104). Extracted spacers were then de-replicated using DADA2 v.1.28.0 (86). To infer spacer origins, the non-redundant spacer sequences were mapped against the bacterial reference database.

### Sequencing data, supplementary data, and code availability

All raw sequencing reads listed in Supplementary Table 1 have been deposited in the NCBI Sequence Read Archive (SRA) under BioProject accession PRJNA1419557. Processed and intermediate analysis files are available on Zenodo at DOI: 10.5281/zenodo.18504637. All code used for data processing, analysis, and figure generation is publicly available on GitHub (https://github.com/Freevini/Old_Raclette), with a permanent release archived on Zenodo (DOI: 10.5281/zenodo.18504637).

## Acknowledgments

We gratefully acknowledge the late Daniel Wechsler for his valuable contributions in the early stages of this research. We are very grateful to the Zufferey family, and especially Maurice and Jean-Jacques Zufferey, who allowed us access to the Jules Zufferey collection of ancient cheeses in Grimentz (Valais, Switzerland). VS thanks the entire Moineau lab for the continued feedback on the project.

We acknowledge Enrique Rayo and Gülfirde Akgül for their valuable laboratory work, Johann Marmy for the design of Figure 1, Josy Fellay for precious information about cheesemaking on the alpine dairy of Moiry, Stephan Häsler for his help clarifying antibiotic usage in herding in Switzerland, and John Haldemann for providing information about the production of Raclette du Valais. Sequencing for this publication was conducted by the Functional Genomics Center Zurich.

S.M. acknowledges funding from NSERC (Discovery program). S.M. holds the Tier 1 Canada Research Chair in Bacteriophages (2011–2025). JBP acknowledges funding from the Swedish Research Council (VR; grants 2019-00299 and 2023-01721) under the frame of JPI AMR (EMBARK and SEARCHER; JPIAMR2019-109 and JPIAMR2023-DISTOMOS-016, respectively), the Data-Driven Life Science (DDLS) program supported by the Knut and Alice Wallenberg Foundation (KAW 2020.0239), and the Swedish Foundation for Strategic Research (FFL21-0174).,

