## supplemental_figures for "Microbial community dynamics in a traditional Swiss mountain cheese over 142 years of cheesemaking"

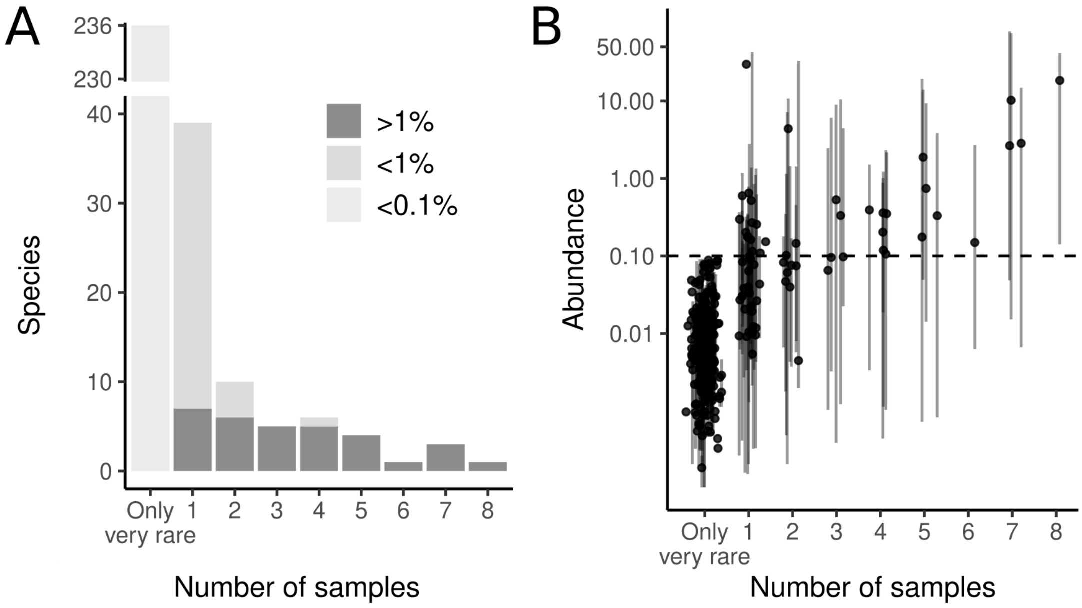


**Supplementary Figure 1.** A) Number of different species detected in the different samples; B) Species prevalence and abundance correlation. The dot is the mean abundance and the line is the spread.

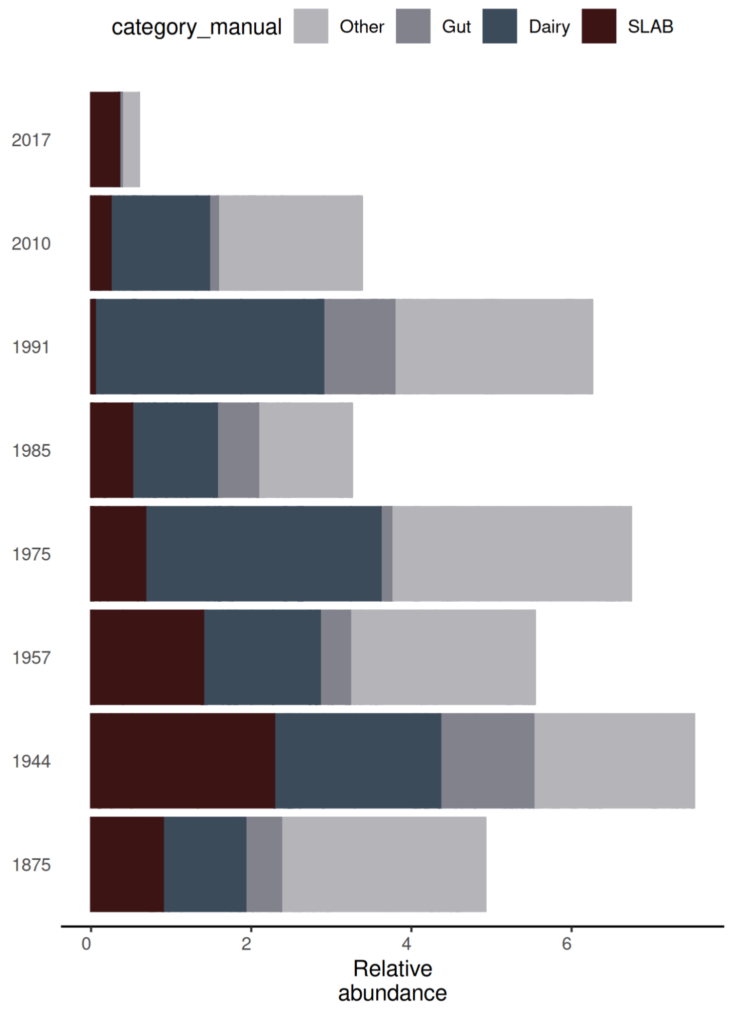


**Supplementary Figure 2.** Relative abundance of the four species types over the different cheese samples.

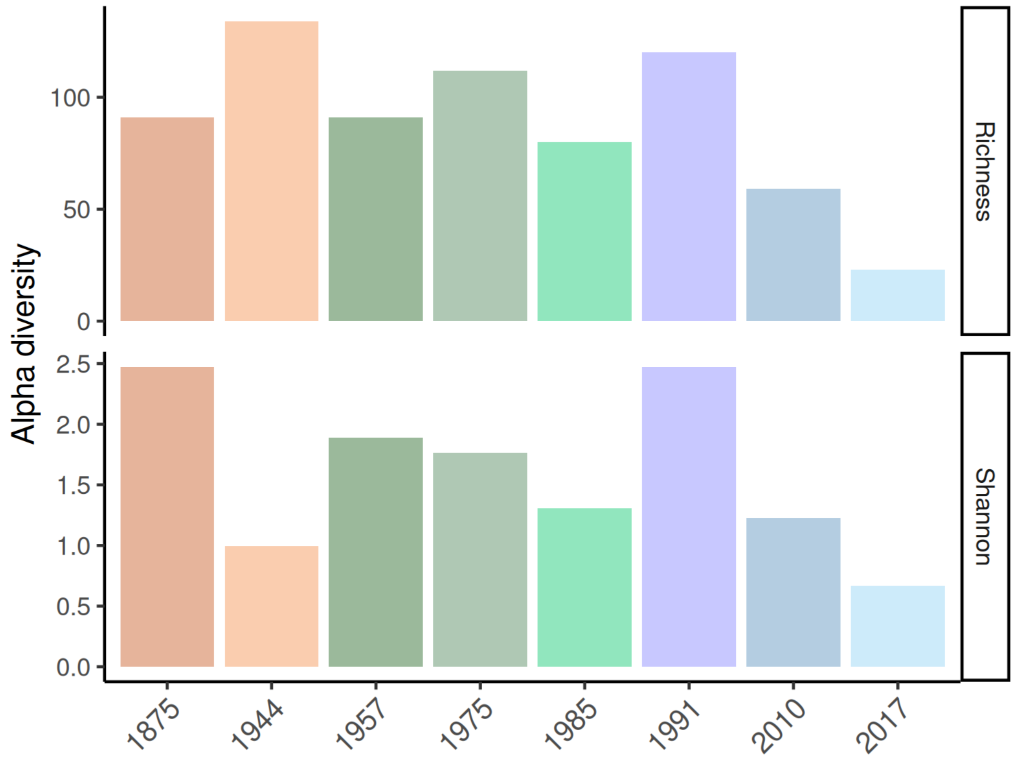


**Supplementary Figure 3.** Species richness and Shannon diversity of bacterial species over time. For both, the linear regression is in both cases not significant.

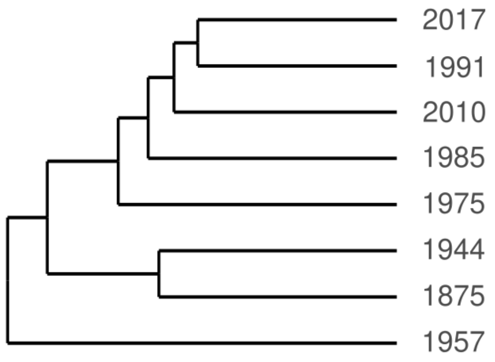


**Supplementary Figure 4.** Dendrogram of samples based on hierarchical clustering of gene family presence/absence.

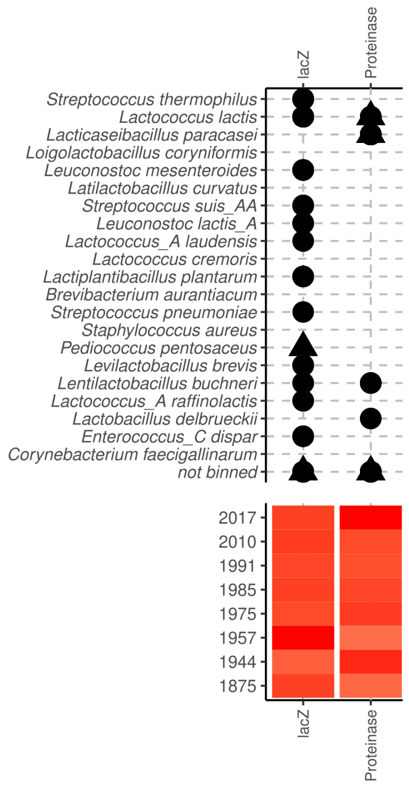


**Supplementary Figure 5.** Cheese phenotypes. Presence of the *lac*-operon in the different MAGs and samples. Genes with the following names were cumulated= *lacZ*, *bgaA*, *lacM*. Right) Presence of the casein-proteinase in the different MAGs and samples. Genes with the following names were cumulated= *prtP*, *prtH*, *prtS*, *prtB*, *prtR*.

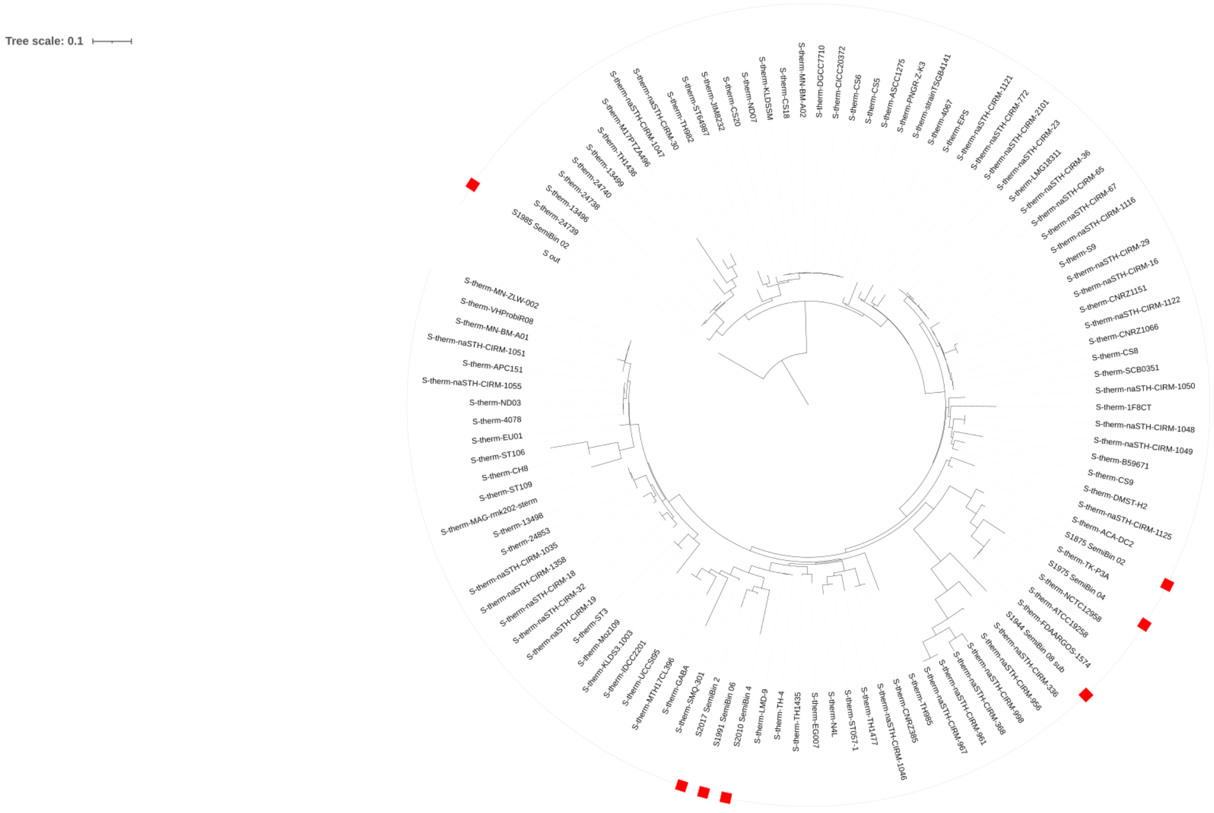

**Supplementary Figure 6.** *Streptococcus thermophilus* phylogeny from Phylophlan of all complete RefSeq genomes and the MAGs generated in this study. The MAGs are highlighted by red squares.

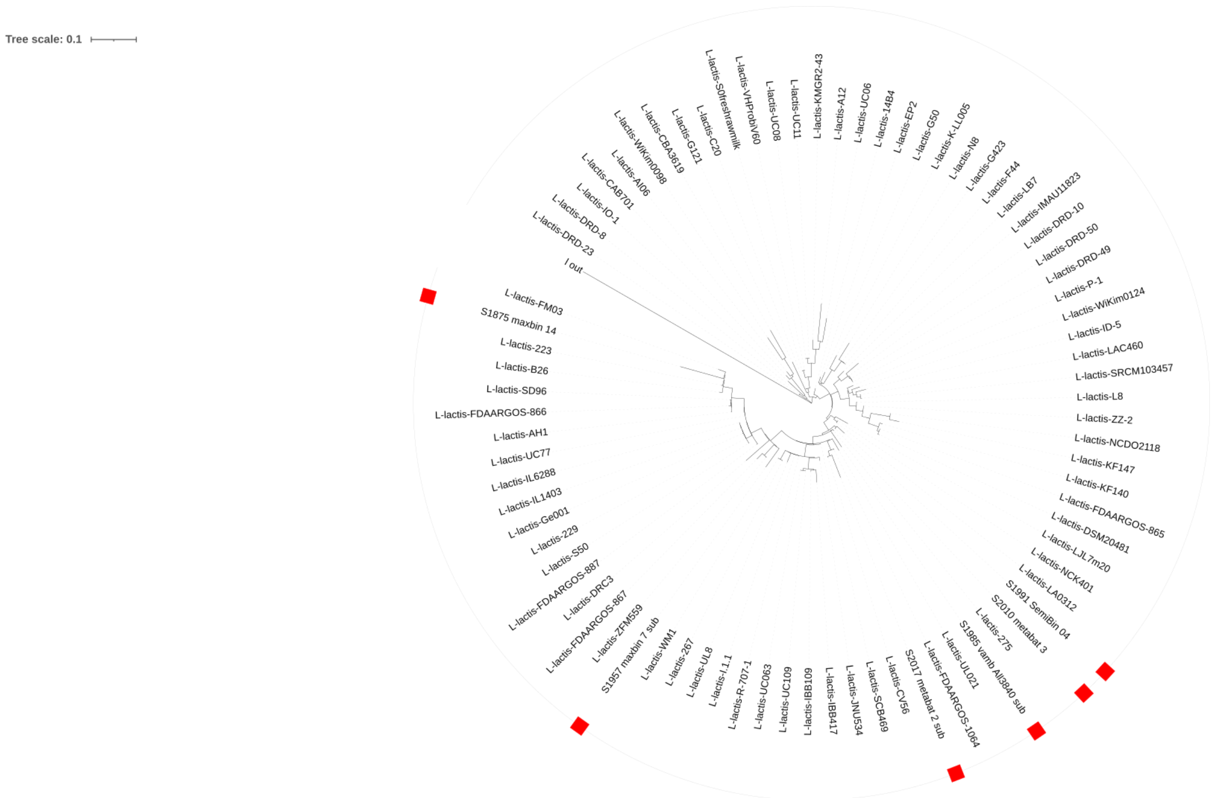


**Supplementary Figure 7.** *L. lactis* phylogeny from Phylophlan of all complete RefSeq genomes and the MAGs generated in this study. MAGs are highlighted by red squares.

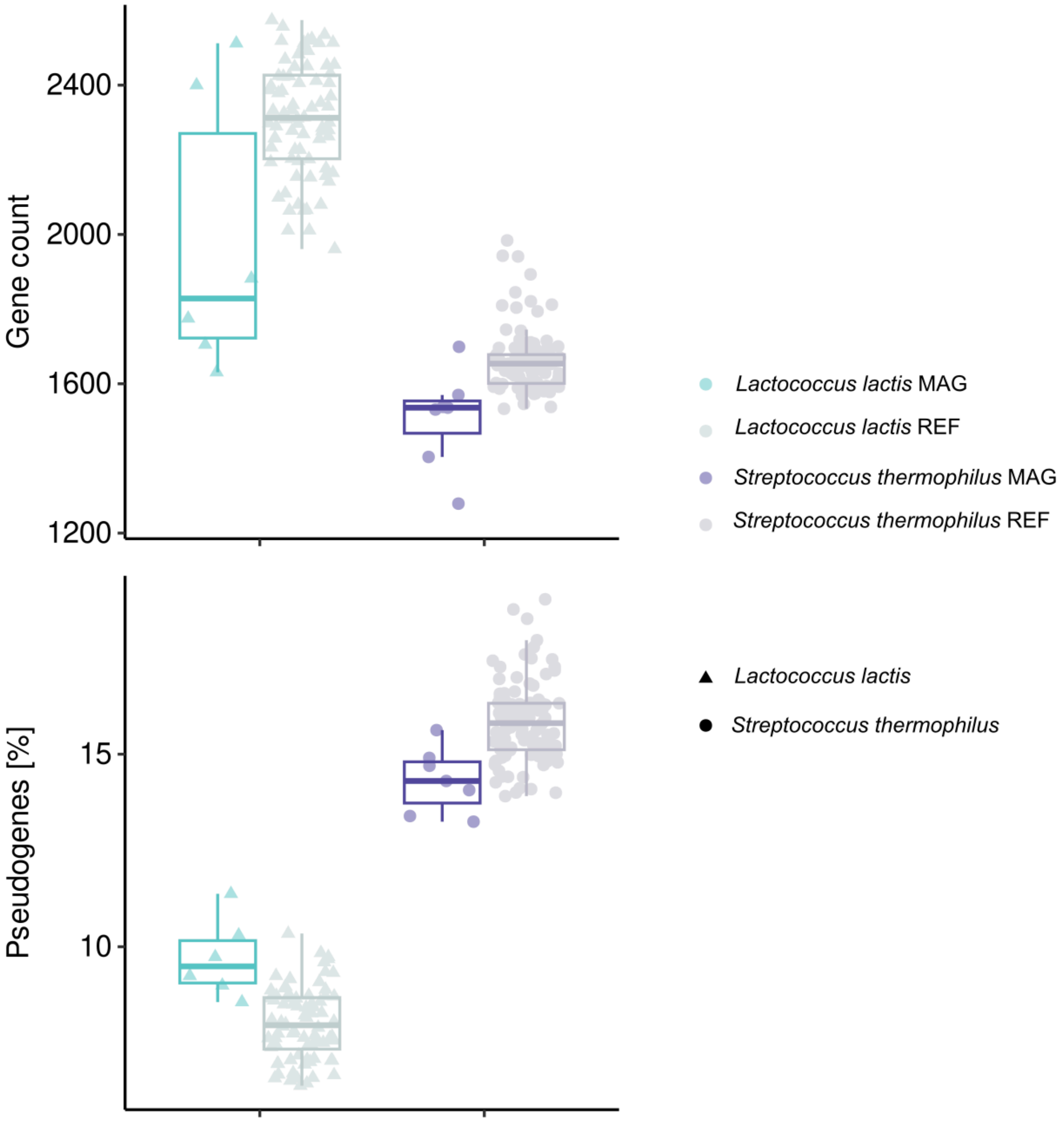


**Supplementary Figure 8.** Number of genes (A) and pseudogenes (B) in reference genomes of *Lactococcus lactis* and *Streptococcus thermophilus* in NCBI (see methods) and in the MAGs of this study. Generally, they are significantly different. This is likely because the MAGs are much more fragmented.

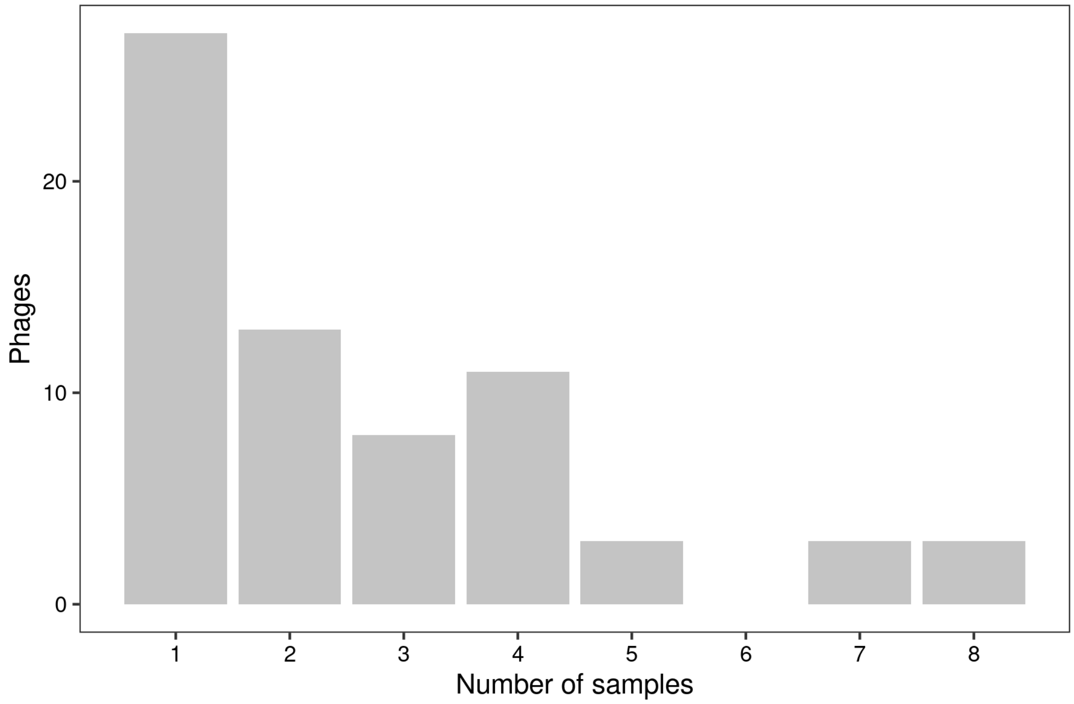


**Supplementary Figure 9.** Occurrence frequency of phages across samples.

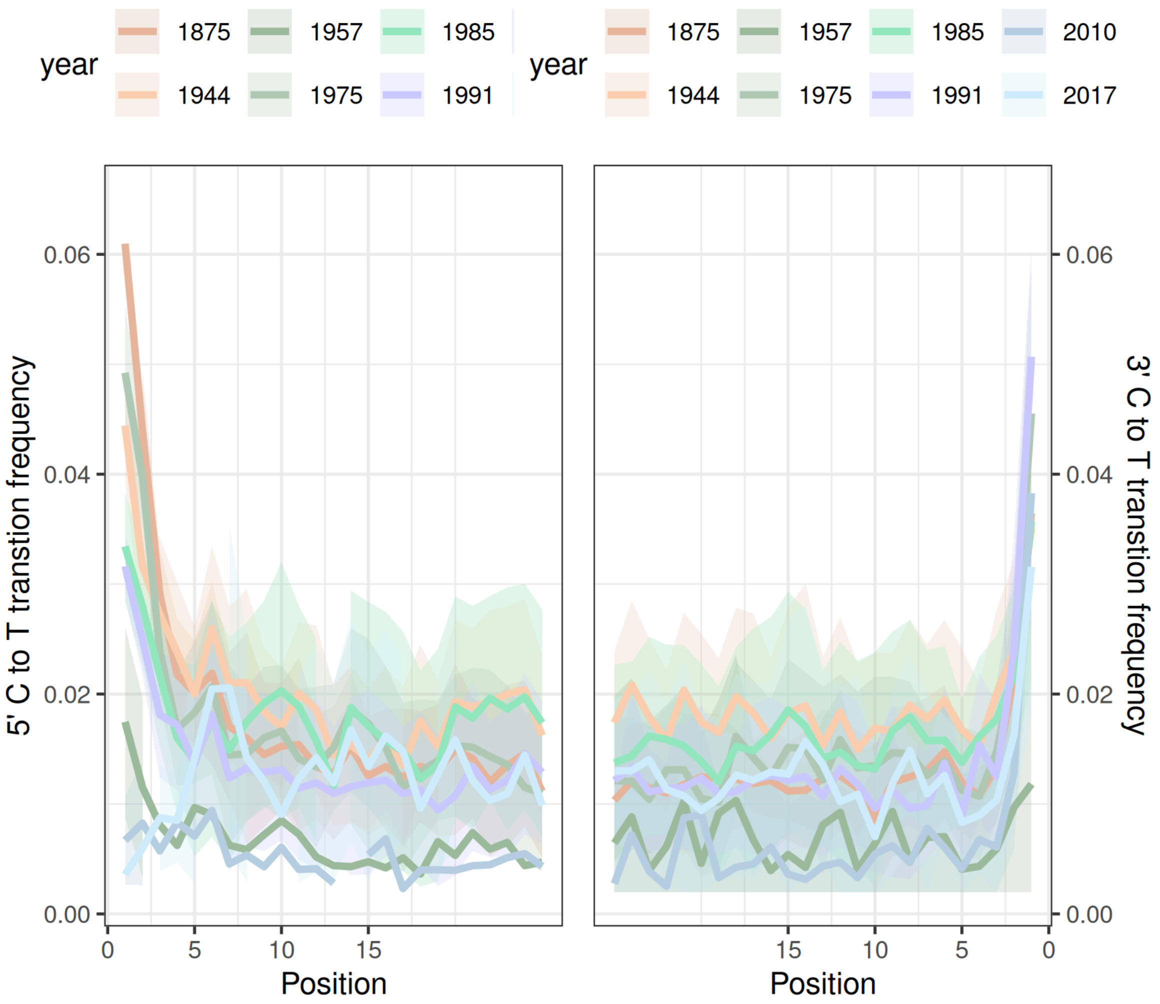


**Supplementary Figure 10.** Damage frequency at the 5′ and 3′ ends of the eight persisting phages across all samples. Older samples consistently show higher terminal damage frequencies than more recent samples, consistent with the ancient origin of these phage sequences.

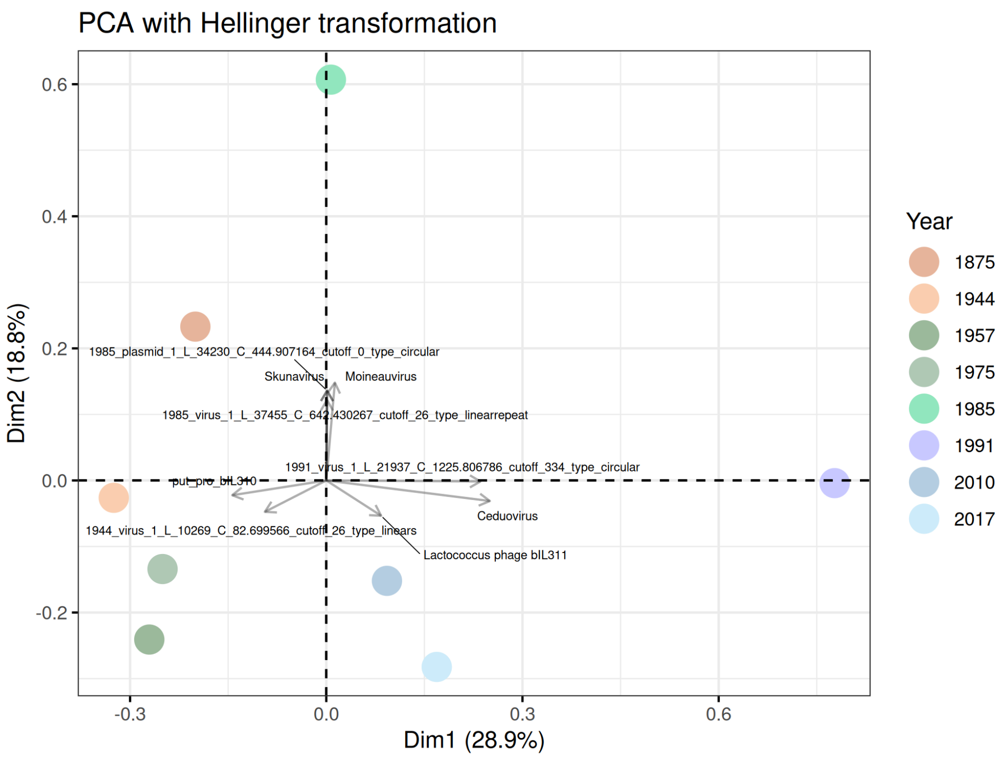


**Supplementary Figure 11.** Principal component analysis PCA with Hellinger transformation based on the mean contig coverage. The significant loadings are illustrated with arrows.

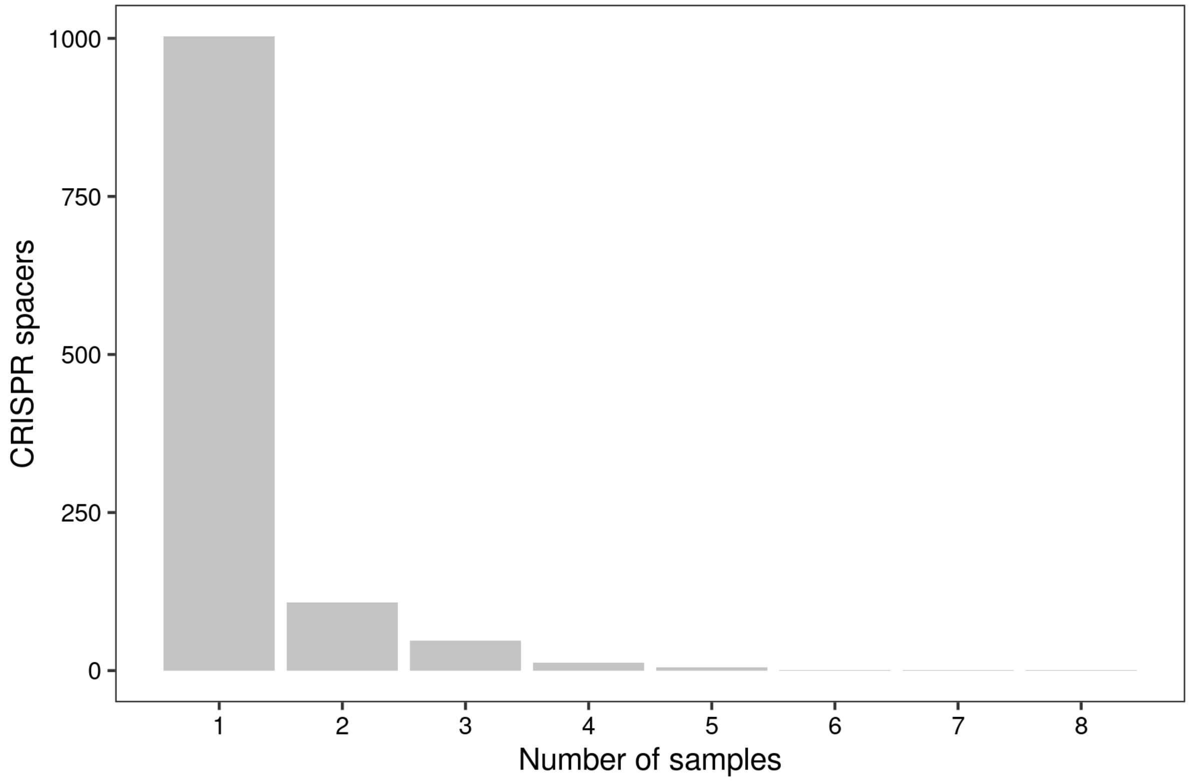


**Supplementary Figure 12.** Number of samples containing the different spacers.

**Table 1.** Raw sequencing data submitted to NCBI bioproject PRJNA1419557.

https://docs.google.com/spreadsheets/d/1HUIden7dyvyN8kxaiLo_j1gQVg_Rvw7PueJqYgvzBcs/edit?gid=0#gid=0
